# Characterization of a novel gene, *srpA*, conferring resistance to streptogramin A, pleuromutilins, and lincosamides in *Streptococcus suis*

**DOI:** 10.1101/2020.08.07.241059

**Authors:** Chaoyang Zhang, Lu Liu, Peng Zhang, Jingpo Cui, Xiaoxia Qin, Lichao Ma, Kun Han, Zhanhui Wang, Shaolin Wang, Shuangyang Ding, Zhangqi Shen

**Affiliations:** Beijing Advanced Innovation Center for Food Nutrition and Human Health, College of Veterinary Medicine, China Agricultural University, Beijing, China; Beijing Key Laboratory of Detection Technology for Animal-Derived Food Safety and Beijing Laboratory for Food Quality and Safety, China Agricultural University, Beijing, China

**Keywords:** SrpA, *Streptococcus suis*, antibiotic resistance, ribosome, ABC-F family proteins

## Abstract

Antimicrobial resistance is undoubtedly one of the greatest global health threats. The emergence of multidrug-resistant (MDR) gram-positive pathogens, like methicillin-resistant *Staphylococcus aureus*, vancomycin-resistant *Enterococcus faecium*, and β-lactamase-resistant *Streptococcus pneumonia*, has severely limited our antibiotic arsenal. Numerous ribosome-targeting antibiotics, especially pleuromutilins, oxazolidinones, and streptogramins, are viewed as promising alternatives against aggressive MDR pathogens. In this study, we identified a new ABC-F family determinant, *srpA*, in *Streptococcus suis* by a comparative analysis of whole genome sequences of tiamulin-resistant and -sensitive bacteria. Functional cloning confirmed that the deduced gene can mediate cross-resistance to pleuromutilins, lincosamides, and streptogramin A in *S. suis* and *S. aureus*. A sequence alignment revealed that *srpA* shares the highest amino acid identity with Vga(E) (36%) and shows canonical characteristics of ABC-F family members. In SrpA–ribosome docked compounds, the extended loop region of SrpA approaches the valnemulin binding pocket in the ribosome peptidyl-transferase center and competes with bound valnemulin. A detailed mutational analysis of the loop residues confirmed that this domain is crucial for SrpA activity, as substitutions or truncations of this region affect the efficiency and specificity of antibiotic resistance. A ribosome binding assay supported the protective effects of SrpA on the ribosome by preventing antibiotic binding as well as displacing bound drugs. These findings clarify the mechanisms underlying resistance to ribosomal antibiotics.

## 1. Introduction

Antimicrobial resistance is now recognized as one of the most serious global threats to human and animal health in the 21st century, and the emergence of multidrug-resistant (MDR) pathogens threatens to cause regression to the pre-antibiotic era [1, 2]. Consequently, infections caused by highly resistant gram-positive organisms, such as staphylococci, streptococci, and enterococci, are considered a major public health issue [3]. In particular, the emergence of methicillin-resistant *Staphylococcus aureus* (MRSA), vancomycin-resistant *Enterococcus faecium* (VRE), and β-lactamase-resistant *Streptococcus pneumonia* have made treatment of the infections very challenging.

More than half of the antibiotics in use target the bacterial ribosome; these inhibitors can paralyze protein synthesis by interacting with the 30S small subunit or 50S large subunit [4-6]. Several ribosome-targeting antibiotics (macrolides, pleuromutilins, lincosamides, streptogramins, and oxazolidinones) are viewed as effective alternatives and last resort drugs for human use (Table S1) [2, 7, 8]. Pleuromutilins were discovered as natural agents in 1951; the C-14 derivatives tiamulin and valnemulin are widely used in veterinary medicine, and retapamulin and lefamulin are approved for human use against MRSA or β-lactamase-resistant streptococci [9, 10]. Structural characterization has revealed that pleuromutilins can bind to the peptidyl-transferase center (PTC) of 23S rRNA, thereby preventing the correct positioning of acceptor or donor substrates at A-sites and P-sites [10-14].

Accumulating evidence suggests that the resistance of gram-positive pathogens to pleuromutilins is increasing; notably, most isolates show cross-resistance to pleuromutilins, lincosamides, and streptogramins A, thereby exhibiting the PLS_A_ phenotype [15-18]. Resistance to pleuromutilins is commonly attributed to the acquisition of endogenous mutations in 23S rRNA or horizontally transmitted resistance determinants, such as the rRNA methyltransferase Cfr or the antibiotic resistance ATP-binding cassette (ABC) F family proteins—Vga/Lsa/Sal/VmlR [4, 7, 19]. Antibiotic resistance ABC-F family genes have been found in the genomes of numerous pathogens and confer resistance to a broad range of clinically relevant antibiotics targeting the ribosomes [20-22]. Unlike other ABC superfamily members that actively pump drugs out of the membranes, the ABC-F family proteins do not fuse to transmembrane domains (TMDs). Accordingly, the mechanism by which ABC-F members mediate antibiotic resistance has been a subject of long-standing controversy. Cryo-electron microscopy (cryo-EM) structures of two ABC-F proteins, MsrE and VmlR, bound to the ribosome, indicate that the protection is conferred by interactions with the antibiotic-stalled ribosomes and displacement of the bound drug [23-25].

*Streptococcus suis* is an emerging zoonotic pathogen; the largest outbreak in humans occurred in China in 2005 (204 infections and 38 deaths) [26]. *S. suis* may also act as a resistance gene reservoir, contributing to the spread of resistance genes to major streptococcal pathogens [27, 28]. During a surveillance study of *S. suis* resistance in China, 42 tiamulin-resistant isolates did not harbor known resistance determinants. In this study, we identified and characterized a novel antibiotic resistance determinant, *srpA (<u>S</u>. suis* <u>R</u>ibosome <u>P</u>rotective <u>A</u>BC-F family protein), including analyses of its physicochemical properties and molecular mechanisms. Our comprehensive analyses, including functional cloning, homology modeling, molecular docking, mutagenic analyses, and ribosome binding assays provide a detailed understanding of the mechanisms by which SrpA confers antibiotic resistance.

## Materials and methods

### 2.1. Bacterial strains

A total of 166 *S. suis* were collected from five provinces in China in 2018, and 72 isolates showed tiamulin resistance. *Escherichia coli* BL21, DH5a, *S. aureus* RN4220, and *S. suis* SD1BY15 were used as hosts for cloning.

### 2.2. Bioinformatics and sequence analysis

The genomic DNA of all tested isolates was extracted as described previously and subjected to whole-genome sequencing [29]. A pan-genome analysis was conducted by the rapid Roary pipeline for 42 resistant candidates (without known determinants) and 94 susceptible strains [30]. The data were processed using the Python package pandas, and ORFs annotated as putative ABC transporter ATP-binding proteins in more than 10 isolates were extracted. The results were visualized using Matplotlib and Seaborn. MEGA7.0 was used to generate a phylogenetic tree of ABC-F family proteins by the neighbor-joining method (Bootstrap: 1000 times) [31]. Sequence logos of corresponding proteins were generated using online tools [32]. The nucleotide sequence of *srpA* and flanking regions has been deposited in the GenBank database under accession number MT550884.

### 2.3. Functional cloning of srpA

The *srpA* gene including 257 bp of its upstream region and 89 bp of its downstream region was cloned from the genomic DNA of HNBY78 (accession number <u>PRJNA616172</u>) and then inserted into the shuttle vector pAM401. The recombinant plasmid was transferred into recipients by electro-transformation.

### 2.4. Homology modeling and molecular docking of SrpA

The 3D model of *srpA* was constructed by homology modeling using the SWISS-MODEL web server (http://www.swissmodel.expasy.org). *srpA* exhibited the highest sequence identity (30%) to the crystal structure of the *Thermus thermophilus* ribosome-MsrE complex (PDB ID: 5Zlu.1.u). Based on the GMQE and QMEAN score, model 1 was selected as the best model [33]. The *S. aureus* ribosome structure was downloaded from the PDB database (5TCU), while the molecular structure of valnemulin was extracted from ChemSpider (http://www.ChemSpider.com). Smina was used as a molecular docking tool to investigate the binding among SrpA, valnemulin, and ribosomes. A total of 2000 docking conformations were extracted by a cluster analysis. A small cluster radius of 10.0 Å for the RMSD cutoff, and a smaller interface cutoff of 10.0 Å, generated better clustering results. The top 10 poses in the largest clusters were obtained and the results were analyzed using PyMol.

### 2.5. Mutational analysis of the extended loop region and ATP hydrolysis glutamates

SrpA mutants were constructed by site-directed mutagenesis using the primers listed in Table S2. A 6×His tag at the N-terminal end of SrpA was introduced in all primers. Sequencing of the complete *srpA* gene from each mutant was conducted to ensure that no extraneous mutations occurred. The expression of SrpA in different derivatives was assessed by western blotting using an anti-His HRP Ab at a 1:2000 dilution.

### 2.6. Ribosome binding assays

The expression and purification of native *srpA* and its mutants were performed as described previously; in brief, *srpA* was ligated into pET28a and transformed to *E. coli* BL21 (DE3) cells [29]. *E. coli* BL21 containing *srpA* were cultured overnight, followed by large-scale induction by IPTG [23]. SDS-PAGE was used to assess the expression of srpA. Proteins were then purified on Ni2+ -NTA affinity resin following a modified version of the manufacturer’s instructions (QIAGEN) [25]. Ribosomes from *S. aureus* RN4220 were purified by differential centrifugation as described by Wu et al. [34]. The cells were resuspended in 4 ml of buffer A (10 mM Tris-HCl containing 4 mM MgCl_2_, 100 mM KCl, and 10 mM NH_4_Cl, pH 7.2). Cell debris was discarded by centrifugation at 30,000 × *g* for 15 min at 4°C. The supernatant was centrifuged at 100,000 × *g* for 120 min at 4°C to pellet the ribosomes [25].

The fluorescent conjugates (tracer) of valnemulin (VAL) and enrofloxacin (ENR) were prepared as described previously [35]. The concentrations of SrpA and ribosomes were optimized for fluorescence polarization immunoassay (FPIA), and the results are depicted in Fig. S*4*.

Concentrations of 1000 nM ribosomes and 5 μM SrpA were chosen for further analyses. The antibiotic–ribosome binding assay was performed by the addition of 70 μL of VAL–DTAF or ENR-AF to purified ribosomes (1000 nM) in 70 μL of buffer A (10 mM Tris [pH 7.5], 60 mM KCl, 10 mM NH4Cl, 300 mM NaCl, 6 mM MgCl_2_, 0.1 mM ATP) at 37°C for 30 min. Reaction mixtures were then shaken for 10 s in the microplate reader, and the FP values for the complex were measured at λex = 485 nm, λ_em_ = 530 nm, cutoff = 515 nm, and G factor = 1.0, while the blank control only contained buffer A, with the same assay conditions [36]. The ability of SrpA to prevent the binding of VAL-tracer to the ribosome was assessed as follows. The ribosomes were preincubated in 70 μL of reaction mixture with 5 μM native SrpA, SrpA mutants, or BSA at 37°C for 30 min before the addition of VAL-tracers. In competitive experiments, the ribosomes were pretreated with VAL-tracers for 30 min and then incubated with native SrpA.

### 2.7. Determination of the genomic locations of srpA

S1-PFGE and Southern blotting were performed to determine the location of *srpA* in *S. suis*. The gel plug of ***srpA*** -positive isolates was prepared as described above and subjected to electrophoretic migration for 20 h [37].

The gel was inverted and placed on the gel box, and a nylon membrane, three dry filter papers, and six inches of cut paper towels were placed on the top of the gel for 16 h. The *srpA* probe was obtained by PCR amplification from the genomic DNA of *S. suis* HNBY78, and then labelled using the DIG High Prime DNA Labelling and Detection Starter Kit I (Roche).

## 3. Results

### 3.1. Identification and characterization of a novel pleuromutilin resistance determinant in S. suis

During routine surveillance of *S. suis* resistance profiles in China (2017 to 2018), 166 isolates were obtained from five provinces, including 94 tiamulin-susceptible isolates and 72 resistant isolates (64– 128 μg/mL). Interestingly, whole-genome sequencing analysis suggested the presence of the PLS_A_ resistance gene, *lsa*(E), in 30 tiamulin-resistant isolates. However, no known resistance determinants were detected in the other 42 strains. Hence, these strains were chosen for an in-depth analysis of the molecular basis of pleuromutilin resistance. A pan-genome analysis was used to compare gene distributions between tiamulin-susceptible and -resistant isolates, with a focus on ORFs annotated as ATP-binding proteins, which have been implicated in pleuromutilin resistance [20]. As depicted in heatmaps (Fig. 1), 68 putative ATP-binding proteins were observed in all annotated isolates. Most ORF_S_ showed no significant correlation with pleuromutilin resistance. However, an ORF annotated as “expZ” was only detected in resistant isolates. We then manually inspected the “expZ” ORFs using the RAST-server and confirmed a 1,368-bp ORF in all 42 resistant isolates. A sequence alignment demonstrated this protein exhibited ∼36% amino acid sequence identity with another antibiotic resistance ABC-F protein Vga(E) and ∼31.6% with VmlR (expZ) [38]. To confirm the role of this putative resistance gene, a 1,980-bp DNA fragment including this ORF and its flanking region was ligated into the shuttle vector pAM401, and the recombinant plasmid was electroporated into *S. suis* SD 1B15 and *S. aureus* RN4220. In comparison with the recipient carrying an empty plasmid, transformants carrying the putative determinant showed significantly higher minimum inhibitory concentrations (MIC) of valnemulin, tiamulin, retapamulin, lefamulin, virginiamycin M, and lincomycin (Table 1), while no differences in MIC were detected for vancomycin, erythromycin, kanamycin, gentamicin, tetracycline, linezolid, and florfenicol. These results suggested that the novel 1,368-bp ORF is a new resistance determinant able to mediate cross-resistance to pleuromutilins, lincosamides, and streptogramin A. Accordingly, we named this ORF SrpA *(<u>S</u>. suis* <u>R</u>ibosome <u>P</u>rotective <u>A</u>BC-F family protein) (accession number MT550884).

**Fig. 1.**
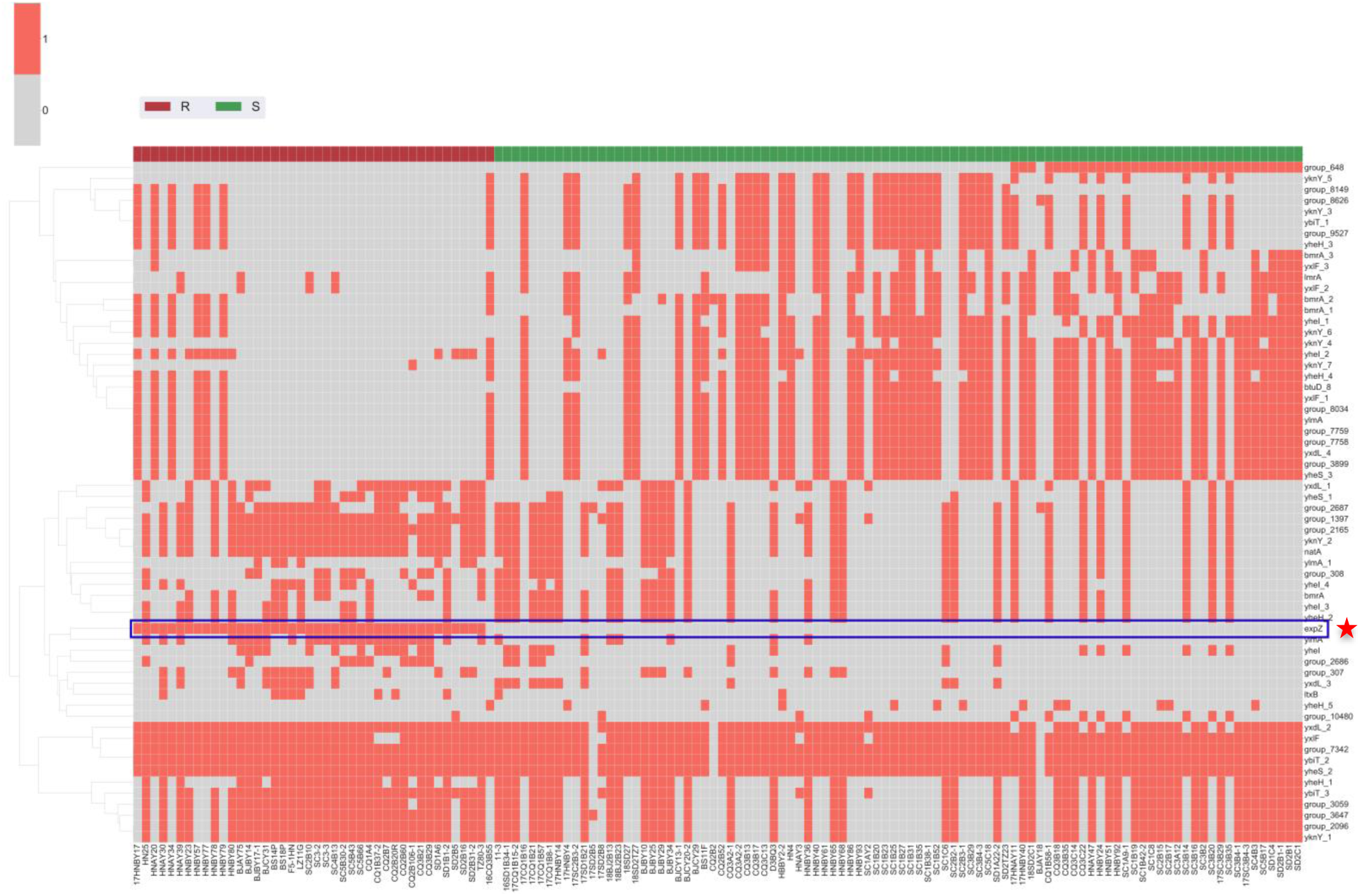
Distribution of putative ATP-binding proteins in tiamulin-resistant and -susceptible isolates. Putative ATP-binding proteins are shown along the ordinate. The names of 136 *S. suis* included in this work are shown on the abscissa. Tiamulin-resistant isolates are shown in red, while susceptible isolates are marked in green. The presence or absence of ATP-binding proteins is indicated by filled or empty squares, respectively. The row “expZ” is marked with a rectangular border and an asterisk.

### 3.2. Relatedness of SrpA with other antibiotic resistance ABC-F proteins

A SMART (http://smart.embl-heidelberg.de/) sequence analysis predicted that SrpA has the canonical architecture of ABC-F family proteins, with two Walker A motifs (residues 36–44 and 303–311), two Walker B motifs (residues 98–102 and 403–407), two ABC signatures (residues 78– 83 and 383–388), two H-loop switches (residues 132–437), and a lack of a transmembrane-associated domain (Fig. S1). Furthermore, for the catalytic motifs of SrpA, the consensus signature sequence was LSGGE, instead of LSGGQ in most ABC superfamily proteins [39].

Thereafter, we built a neighbor-joining tree to estimate the relationships of SrpA with other antibiotic resistance ABC-F family proteins. As depicted in Fig. 2, the phylogenetic tree was characterized by five deep-branching lineages. SrpA belonged to a lineage including Vga homologs, Msr homologs, VmlR and Lmo0919. A sequence alignment showed that SrpA shares over 30% amino acid identity with loci in this lineage, and the closest homologue was Vga(E) in *S. aureus* (36%). Although antibiotic resistance ABC-F family proteins are found in a variety of bacterial groups, SrpA is the first antibiotic resistance ABC-F protein found in *S. suis*.

**Fig. 2.**
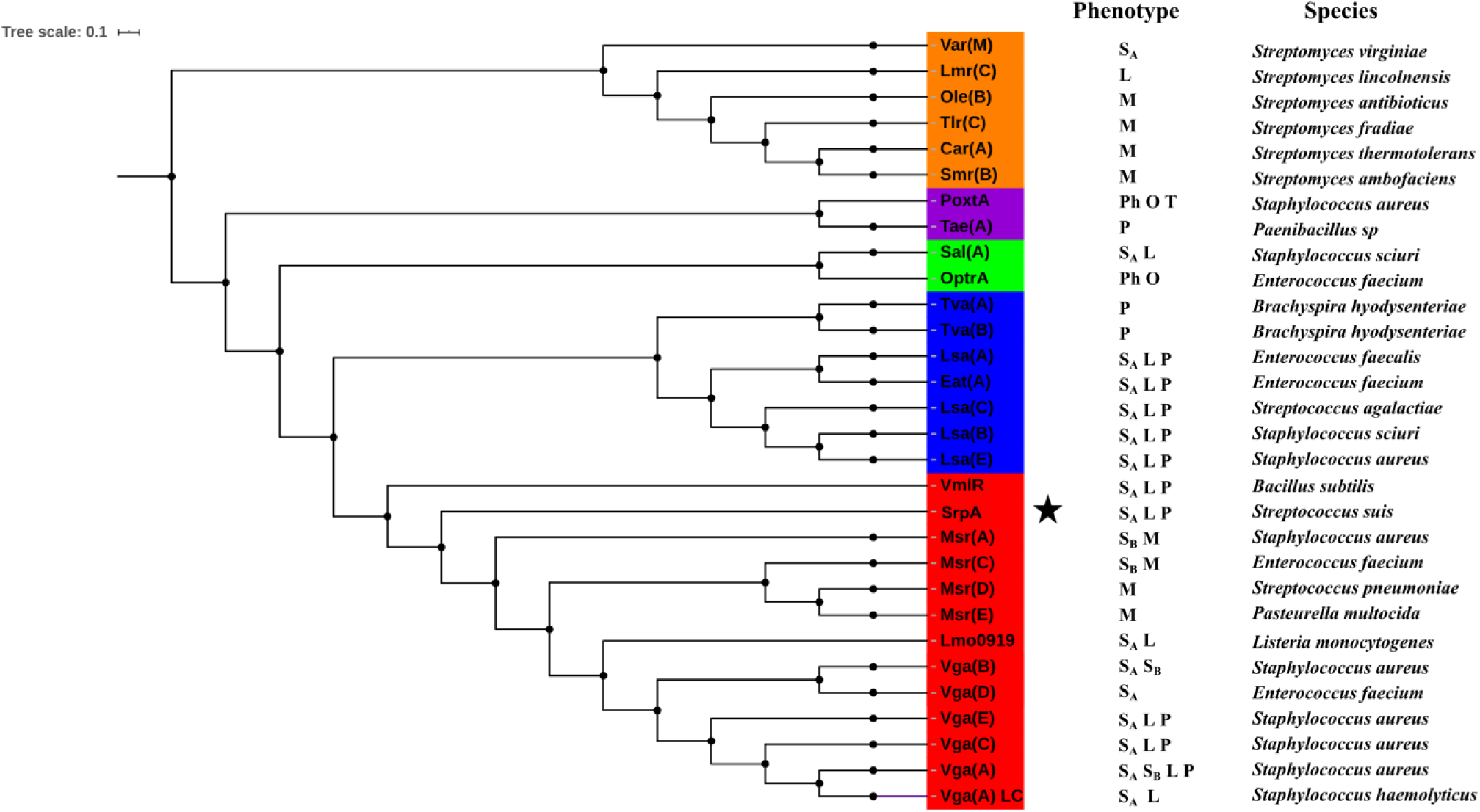
Phylogenetic tree of antibiotic resistance ABC-F proteins constructed using the neighbor-joining method. The tree represents the consensus obtained after 1000 replicates. Amino acid sequences were extracted from NCBI. Antibiotics affected by the different proteins are indicated and the species in which the various antibiotic resistance ABC-F have been described for the first time are also indicated. S_A_, streptogramin A; S_B,_ streptogramin B; P, pleuromutilins; L, lincosamides; M, macrolides; Ph, phenicols; O, oxazolidinones; T, tetracyclines. The novel antibiotic resistance ABC-F determinant SrpA is indicated by an asterisk.

### 3.3. Homology modeling and molecular docking of SrpA

A 3D model of SrpA was created using the SWISS-MODEL tool, and the crystal structure of Msr(E) (PDB ID: 5Zlu.1.u), which shared 30% sequence identity with SrpA, was selected as the most suitable template. The structure of SrpA was predicted to possess conserved features of antibiotic resistance ABC-F family proteins, such as two ABC transporter domains (ABC1 and ABC2) carrying highly conserved nucleotide-binding sites (NBSs) (residues 36–43 and 303–310) assembled at its base, and a domain linker (residues 163–250) that forms two long crossed helices (α1 and α2) connected by an extended loop (residues 203–224) (Fig. 3*A*) [20, 25].

**Fig. 3.**
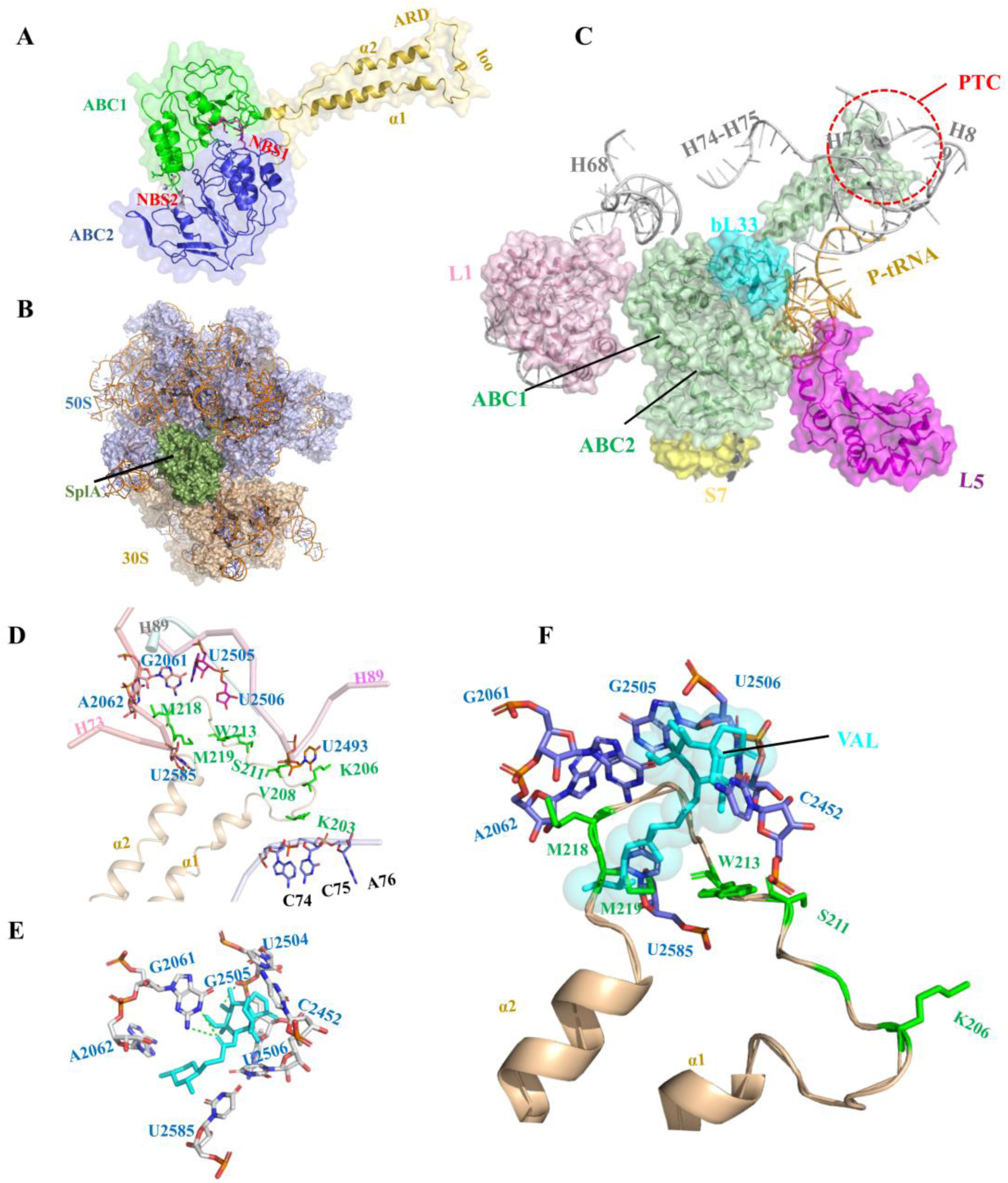
Homology modeling and molecular docking of SrpA. (A) Three-dimensional (3D) structures of SrpA generated by the SWISS-MODEL server; protein shown in cartoon with secondary structure elements of ABC1 (green), ABC2 (blue), and domain linker (yellow orange) labeled. (B) Binding mode of SrpA (green), and ribosomes (50S slate, 30S light orange). (C) Interactions between the SrpA (green) ABC1 domain and the 50S subunit r-protein L1 (pink), bL33 (cyan), 23S rRNA helices H68, H73–H75, H89 (white) and between the ABC2 domain and the 30S subunit r-protein S7 (yellow), r-protein L5 (purple), and tRNA (wheat). (D) Orientation of the SrpA extended loop (yellow orange) and surrounding key PTC residues. (E) Valnemulin-binding pocket (cyan) in the valnemulin–ribosome complex. (F) Superposition of ribosome-bound valnemulin onto the SrpA–ribosome complex.

Several studies have demonstrated that antibiotic resistance ABC-F proteins bind to the ribosome and allosterically dissociate the drug from its binding site [23, 24]. In the present work, molecular docking was used to analyze SrpA–ribosome interactions. The crystal structures of the *S. aureus* 70S ribosome were extracted from the PDB database (ID: 5TCU), and the complex was docked with a 6° Euler angle sampling size using ZDOCK 3.0.2 (Fig. 3*B*). As shown in Fig. 3*C*, the ABC1 domain of SrpA faces the L1 stalk of the 50S subunit, whereas the ABC2 domain contacts the r-protein L5, S7 and the elbow of P-tRNA. In contrast to the few ribosome contacts observed for the two ABC domains, the elongated domain linker established extensive contacts with ribosomes, as it stretched in parallel with H74–H75 of 23S rRNA and the acceptor arm of P-site tRNA. In addition, the tip loop region was inserted deep into the PTC, approaching the 23S rRNA helices H89 and H73. The residues Lys203, Arg204, and Lys205 in the loop region directly contacted the CCA end of a P-site tRNA, while the Lys206–Ser224 residues and their side chains interacted with the flexible nucleotides U2585, A2062, G2061, U2505, U2506, C2452, U2492, and U2493 (Fig. 3*D*).

Interestingly, the aforementioned nucleotides seemed to constitute the valnemulin-binding pocket, as the valnemulin-ribosome docked compounds demonstrated that G2061 and U2505 can hold the two sides of valnemulin by hydrogen bonds, while the other residues (e.g., U2585, A2062, U2506, and C2452) were also located in close proximity to valnemulin (Fig. 3*E*). Superimposing the SrpA-ribosome complex and valnemulin-ribosome complex revealed steric overlap between the SrpA loop region and valnemulin in the valnemulin-binding pocket; in particular, the Met219 residue directly clashed with the C14 extensions of valnemulin (Fig. 3*F*).

### 3.4. Mutagenic analyses of the ATP hydrolysis site and extended loop region

ATP hydrolysis appears to play a key role in the functioning of ABC-F proteins, as ATP hydrolysis-deficient EttA could not be released from ribosomes, and ATP hydrolysis-deficient MsrE and Vga(A) show reduced or abolished antibiotic resistance [23, 39, 40]. In this work, the conserved catalytic glutamates following the Walker B motifs in both NBDs (Glu103 in NBD1 and Glu in 408 NBD2) were mutated to glutamine. A growth curve showed that this ATP hydrolysis-deficient SrpA mutant (E103Q/E408Q) did not significantly affect the cell growth of *S. aureus* (Fig. S3). However, the antibiotic resistance of the SrpA 103Q/E408Q mutant was significantly lower than that of the wild type, indicating that ATP hydrolysis was crucial for PLS_A_ resistance (Table 1).

**Table.1.**
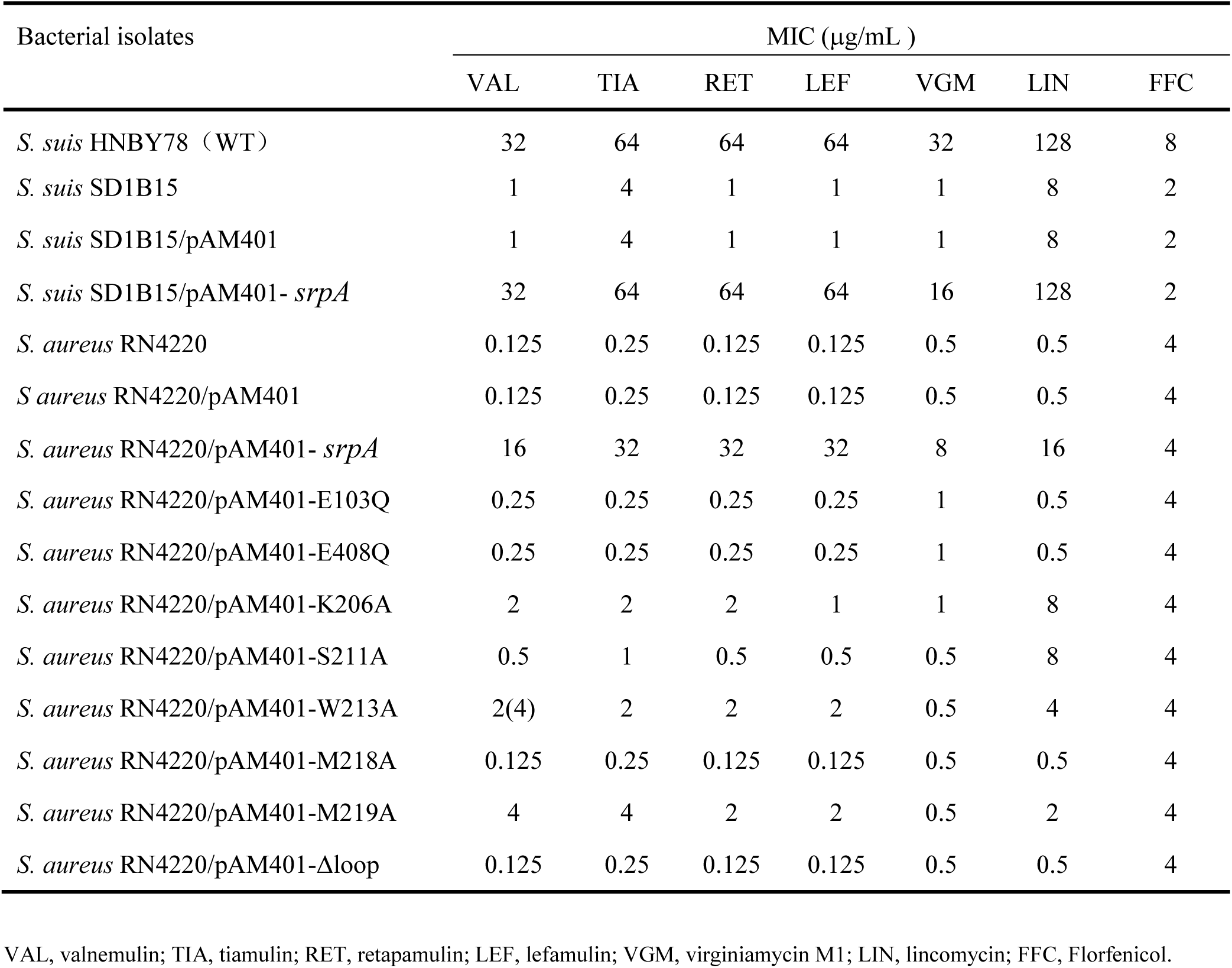
Minimum inhibitory concentrations of tested antimicrobial agents for the studied bacterial isolates

| Bacterial isolates | MIC (µg/mL ) |  |  |  |  |  |  |
| --- | --- | --- | --- | --- | --- | --- | --- |
|  | VAL | TIA | RET | LEF | VGM | LIN | FFC |
| *S. suis* HNBY78 (WT) | 32 | 64 | 64 | 64 | 32 | 128 | 8 |
| *S. suis* SD1B15 | 1 | 4 | 1 | 1 | 1 | 8 | 2 |
| *S. suis* SD1B15/pAM401 | 1 | 4 | 1 | 1 | 1 | 8 | 2 |
| *S. suis* SD1B15/pAM401- *srpA* | 32 | 64 | 64 | 64 | 16 | 128 | 2 |
| *S. aureus* RN4220 | 0.125 | 0.25 | 0.125 | 0.125 | 0.5 | 0.5 | 4 |
| *S. aureus* RN4220/pAM401 | 0.125 | 0.25 | 0.125 | 0.125 | 0.5 | 0.5 | 4 |
| *S. aureus* RN4220/pAM401- *srpA* | 16 | 32 | 32 | 32 | 8 | 16 | 4 |
| *S. aureus* RN4220/pAM401-E103Q | 0.25 | 0.25 | 0.25 | 0.25 | 1 | 0.5 | 4 |
| *S. aureus* RN4220/pAM401-E408Q | 0.25 | 0.25 | 0.25 | 0.25 | 1 | 0.5 | 4 |
| *S. aureus* RN4220/pAM401-K206A | 2 | 2 | 2 | 1 | 1 | 8 | 4 |
| *S. aureus* RN4220/pAM401-S211A | 0.5 | 1 | 0.5 | 0.5 | 0.5 | 8 | 4 |
| *S. aureus* RN4220/pAM401-W213A | 2(4) | 2 | 2 | 2 | 0.5 | 4 | 4 |
| *S. aureus* RN4220/pAM401-M218A | 0.125 | 0.25 | 0.125 | 0.125 | 0.5 | 0.5 | 4 |
| *S. aureus* RN4220/pAM401-M219A | 4 | 4 | 2 | 2 | 0.5 | 2 | 4 |
| *S. aureus* RN4220/pAM401-Δloop | 0.125 | 0.25 | 0.125 | 0.125 | 0.5 | 0.5 | 4 |

VAL, valnemulin; TIA, tiamulin; RET, retapamulin; LEF, lefamulin; VGM, virginiamycin M1; LIN, lincomycin; FFC, Florfenicol.

The extended loop region of antibiotic resistance ABC-F proteins are collectively referred to as the antibiotic resistance domain (ARD), as the tip loop region accesses the PTC of the ribosome and directly encroaches the binding site of PTC-targeting antibiotics [21, 41]. Based on previous studies and the insights gain from the docked compounds, the loop region of SrpA appears to play a major role in the efficiency and specificity of antibiotic resistance.

Based on multiple sequence alignment of SrpA, Vga homologs, Msr homologs, VmlR, and Lmo0919 obtained using ClustalX, this short stretch corresponded to residues Lys203–Ser224 in SrpA. Consensus sequence logos were generated for the extended loop region (Fig. 4*A*). A total of 4 highly conserved amino acid residues were found in SrpA: Lys203, Lys206, Gly207, and Ala217. The arrangement of amino acids at other sites was polymorphic, especially for residues Val208, Trp213, Met218, Gly220, and Ser221. To further confirm the effect of the extended loop region on the efficiency and specificity of antibiotic resistance, we generated mutations in the SrpA loop region at positions Trp206Ala, Ser211Ala, Trp213Ala, Met218Ala, and Met219Ala. The introduction of point mutations at Lys203, Val208, Gly220, and Ser221 was not successful, as the mutants were quite instable. In addition, recombinant vector pAM401 with the truncation of the extended loop region (residues Lys203–Ser224, Δloop) was also constructed by homologous recombination.

**Figure 4.**
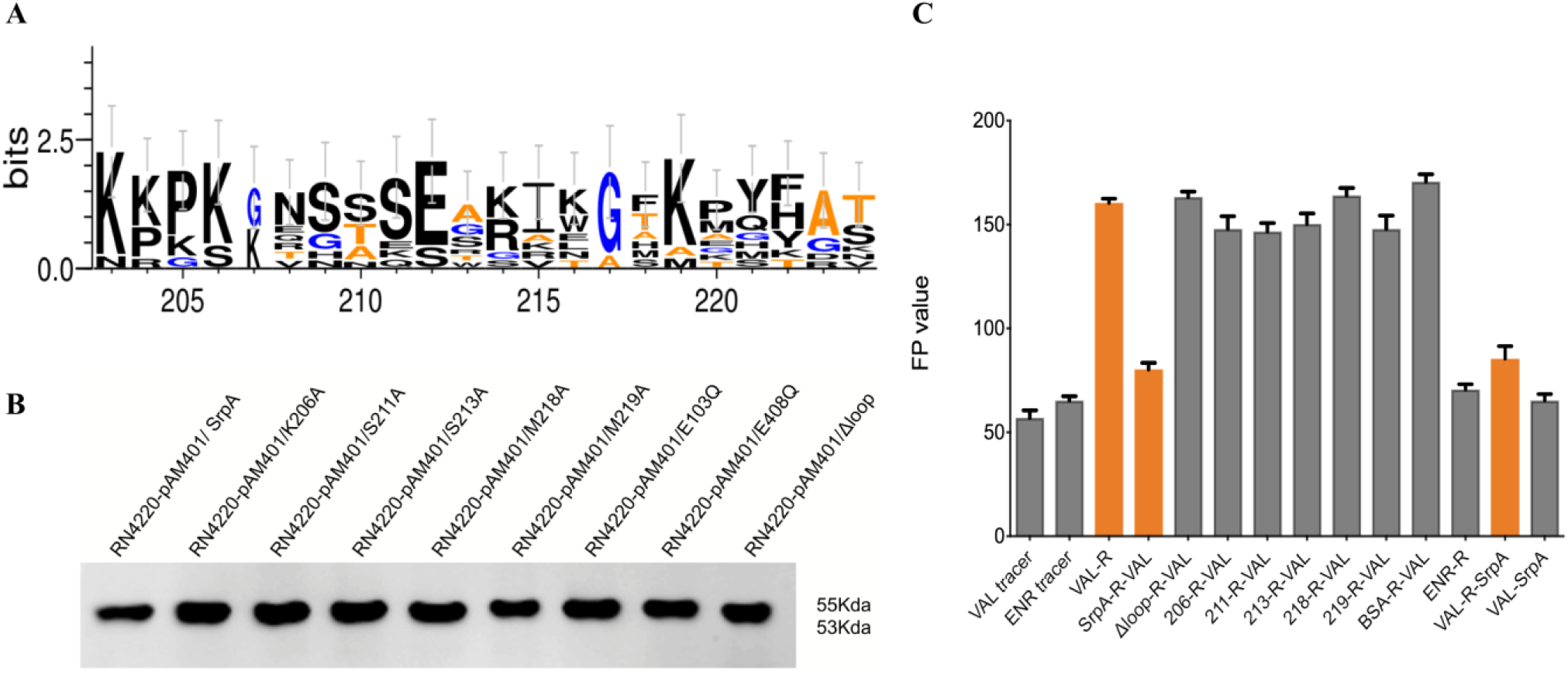
Key amino acids of SrpA for ribosome protective activity. (A) Sequence logos of the loop region amino acid residues generated for SrpA, Vga(A), Vga(E), Msr(E), VmlR, and Lmo0919. The height of each letter is proportional to the frequency of the correspondingly amino acid residues in a given site (B) Western blot analysis of native SrpA and SrpA mutants in *S. aureus* RN4220. (C) Ribosome binding assay (Column 1 and 2 show the FP values of VAL and ENR tracer. In column 3, 1000 nM purified *S. aureus* ribosomes added to the system, while in column 4, the ribosomes were preincubated with 0.5 μM SrpA for 30 min. In columns 5–11, SrpA was replaced with mutants (K206A, S211A, W213A, M218A, M219A, Δ(K203-S224)) or BSA. In column 12, 1000 nM purified *S. aureus* ribosomes was added to the ENR-tracer system. In column 13, the purified *S. aureus* ribosomes were first pretreated with VAL-tracer for 30 min and 5 μM SrpA was introduced to the complex. In column 14, 5 μM SrpA was added directly to the VAL tracer without ribosomes. Results are means of three independent repeats; error bars represent SDs. No ribosome addition was included as a control. R, *S. aureus* ribosomes.

An antibiotic susceptibility test showed that mutations in residues in the loop domain indeed influenced the PLS_A_ resistance phenotype in vivo (Table 1). Compared with isolates harboring intact SrpA, strains bearing mutations at residues 206, 211, 213, 218, and 219 showed a 2- or 4-fold decrease in lincomycin resistance and an 8- to 16-fold decrease in the MIC of virginiamycin. For pleuromutilin compounds, strains carrying the Trp206Ala, Trp213Ala or Met219Ala substitutions exhibited 2- to 16-fold decreases in the MICs of valnemulin, tiamulin, retapamulin, and lefamulin. The Ser211Ala substitution resulted in 16- to 32-fold lower MICs. Intriguingly, the truncation of the extended loop region (residues Lys203–Ser224, Δloop) or even the mutation of a single residue (Met218Ala) could completely abolish the ability to mediate PLS_A_ resistance. As a control, the florfenicol resistance of all derivates remained unchanged. Taken together, these results suggested that the extended loop region, particularly Met218, was crucial for PLS_A_ resistance in vivo.

A western blot analysis with an anti-6xHis antibody was used to examine the expression of SrpA derivatives in the transformants. A single protein band with a molecular weight of about 55 KDa was detected in positive control *S. aureus* RN4220 harboring native SrpA and its seven mutant derivatives (Trp206Ala, Ser211Ala, Trp213Ala, Met218Ala, Met219Ala, Glu103Gln, and Glu408Gln). The transformant carrying the *srpA*-Δloop migrated shows a protein size of 53 kDa (Fig. 4*B*); this size decrease might be explained by the deletion of the extended loop region.

### 3.5. Ribosome binding assays

To further elucidate the mechanism of action of *srpA*, a modified FPIA was used to measure the interaction between SrpA and ribosomes. The fluorescence polarization (FP) value is determined by the size of fluorescent-labeled antibiotics tracer. For an unbound tracer with a small size and fast Brownian rotation, the FP value is low, whereas for the tracer–ribosome complex, the FP value is high. The initial FP values of the tracers VAL-DTAF and ENR-AF were 58 and 66, respectively (Figure 4*C*). After the introduction of 1000 nM purified *S. aureus* ribosomes, the FP value of the Val-ribosome mixture rose to 160. For the ENR mixture, the FP value showed nearly no variation, suggesting specific binding between valnemulin and bacterial ribosomes.

Interestingly, when *S. aureus* ribosomes were preincubated with SrpA (5 µM) for 30 min and subsequently titrated into VAL-DTAF, the FP value of the complex decreased to approximately 79, far less than the value for the direct VAL-ribosome interaction, revealing that SrpA can interact with ribosomes and prevent the subsequent binding of VAL-tracer. When the ribosome was pretreated with VAL-tracer for 30 min and the complex was incubated with SrpA (5 µM) for another 30 min, the FP value was nearly 88, indicating that SrpA displaces prebound VAL-tracer from ribosomes. When *S. aureus* ribosomes were pretreated with BSA instead of SrpA, the FP value was similar, confirming that the ability to rescue ribosomes from valnemulin binding was a specific property of SrpA. The extended loop region played an important role in mediating antibiotic resistance, and this result was also mirrored by FPIA. The preincubation of ribosomes with the SrpA-Δloop mutants did not reduce the level of ribosomally associated Val tracer. Furthermore, our data also demonstrated the importance of the loop residues Trp206, Ser211, Trp213, Met218, and Met219, as mutations to Ala caused a significant loss of ribosomal protection activity.

### 3.6. Characterization of srpA in S. suis

*srpA*-positive *S. suis* were collected from five provinces separated by great geographical distance, indicating the wide distribution of this gene in China. S1-PFGE and Southern blotting revealed that *srpA* was located in the chromosomal DNA of all positive isolates (Figure S5). Isolates were subdivided into two types according to genetic context. The type I synteny was identical in 40 isolates, with a 2,078-bp penicillin binding protein gene *pbp2b* located immediately downstream of the *srpA* gene and an 804-bp MBL fold metallo-hydrolase gene *yycJ* upstream of *srpA* (Figure 5). Interestingly, these two motifs were highly conserved in *S. suis*, even in isolates without *srpA*, and no transposons or insertion element sequences were identified in the vicinity of *srpA*. Type II isolates showed two directly repeated IS*Ssu8*-like elements (93% identity) in the region flanking *srpA*. Reverse PCR confirmed that the insert elements bracketing *srpA* could recombine and form a 3,195 bp circular intermediate including *srpA* and one copy of the IS*Ssu8*-like locus.

**Figure 5.**
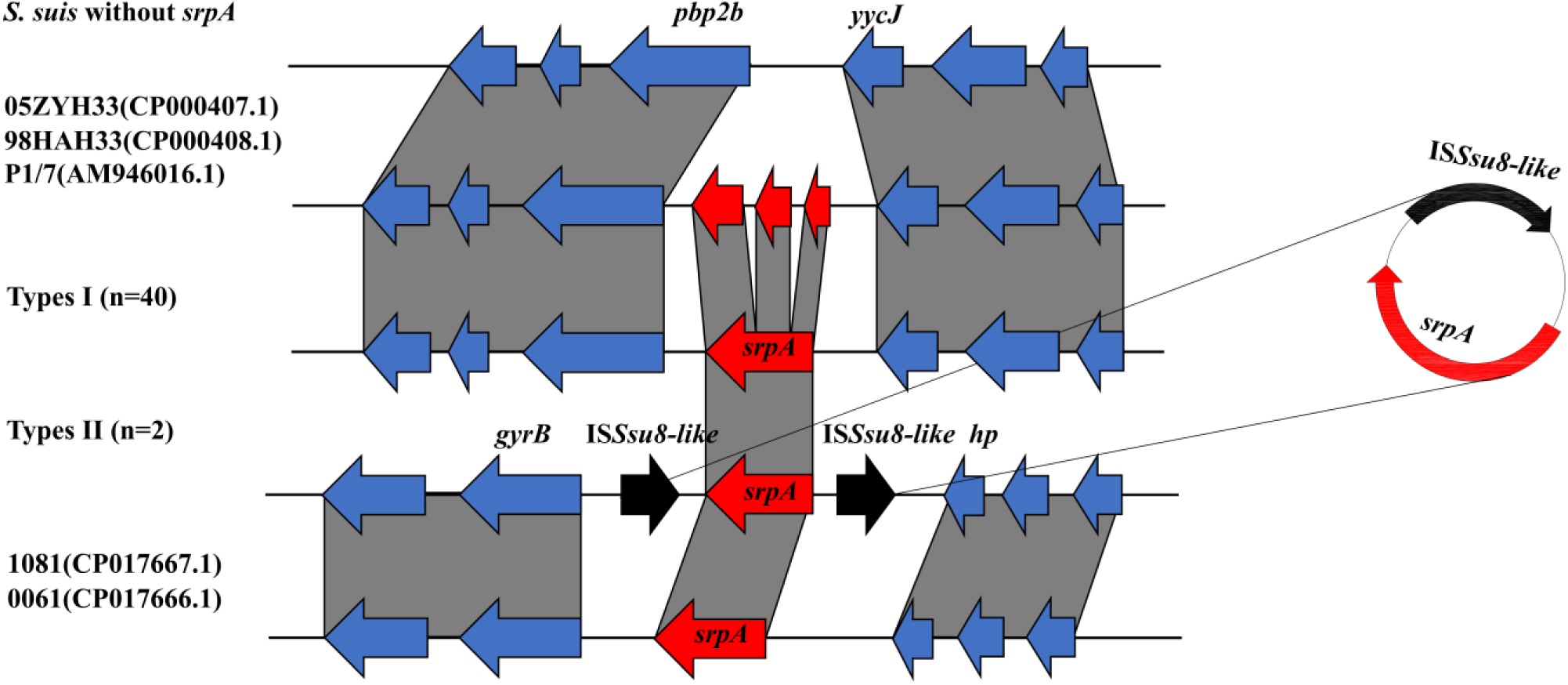
Genetic context of the *srpA* gene. Regions with >90% nucleotide identity are connected by grey zones. *srpA* is shown in red, IS elements in black, and other genes in blue. *S. suis* 05ZYH33(CP000407.1), 98HAH33(CP000408.1), P1/7(AM946016.1), 1081(CP017667.1), and 0061(CP017666.1) were obtained from NCBI.

## 4. Discussion

Since the discovery of penicillin in 1929, antimicrobial agents have been indispensable for the control of bacterial infections in clinical settings and in veterinary medicine [42]. However, their efficacy has been hampered by the emergence of antimicrobial resistance in bacteria. Many infections are caused by highly resistant gram-positive organisms, and the emergence of MRSA, VRE, or β-lactamase-resistant streptococci has severely limited the antibiotic arsenal.

To counteract these MDR bacteria, some ribosome-targeting antibiotics have been developed as alternative or adjuvant agents in combination with β-lactams or vancomycin [3]. Ribosomes are the protein-synthesizing factories of the cell, and more than half of antibiotics used in clinical settings target bacterial the ribosome [6]. Despite their diverse chemical structures, all ribosomal active antibiotics interact with the ribosome PTC or nascent peptide exit tunnel region and possess overlapping binding sites (A-, P-, and E-sites) [4, 7, 12].

*S. suis* is an important pathogen in the pig industry and an agent of zoonosis, transmitted by close contact with infected pigs or pork-derived products [43, 44, 45]. In 1998 and 2005, two large-scale outbreaks of human *S. suis* infections occurred in China, with high morbidity and mortality [46]. During a surveillance study of *S. suis* resistance in China, we found a novel antibiotic resistance ABC-F resistance determinant named *srpA*. The ORF encodes a protein of 461 amino acids and confers cross-resistance to pleuromutilins, lincosamides, and streptogramin A in *S. suis* and *S. aureus*. Remarkably, all *srpA-*positive transformants showed high levels of resistance to the newly approved pleuromutilin derivate lefamulin (2019, FDA). Lefamulin was the first pleuromutilin approved for systemic human use and is regarded as a potential treatment option for MRSA or MDR streptococci; accordingly, lefamulin resistance in *S. aureus* and *S. suis* is an alarming observation. A sequence alignment of SrpA with other antibiotic resistance ABC-F family proteins revealed that SrpA belongs to a lineage containing Vga homologs, Msr homologs, Lmo0919, and VmlR. The tertiary structure of SrpA was more similar to that of MsrE, including two ATP-binding domains connected by an 87-amino acid linker. Furthermore, SrpA lacked the C-terminal extension (CTE) found in VmlR (residues 483–547) and Vga homologs (residues 460–520), potentially explaining the shorter length of SrpA (461 aa) than these other proteins (521–547 aa) [22].

ATPase activity is important for the function of some ABC-F family members. Mutations of the catalytic glutamate residues following the Walker B motif in EttA influence cell growth by inhibiting protein synthesis, while in VgaA_LC_ and LsaA, hydrolytically inactive mutants inhibit the peptidyl transferase activity of the ribosome [22, 39]. In the case of MsrE, the glutamates of both LSGGE motifs are positioned close to the γ-phosphate of AMP-PNP and are likely involved in ATP hydrolysis, while glutamate mutants show reduced ribosome binding in vitro and AZM resistance in vivo [23]. In the present work, the SrpA Walker B (E103/408Q) mutants did not show PLS_A_ resistance, suggesting that ATP hydrolysis is pivotal for SrpA activity.

Molecular docking was used to obtain insight into the SrpA-ribosome interaction. Based on the complex model, ABC1 and ABC2 of SrpA established massive interactions with ribosomal protein and the elbow of the P-tRNA, while the interdomain linker, especially the extended loop region, projected deep into the ribosome PTC, interacted with the CCA arm of P-tRNA, and was located in proximity to 23S rRNA helices H89 and H73. The architecture and conformation of the SrpA-ribosome interactions were very similar to those of MsrE/VmlR, suggesting that the binding site of antibiotic resistance ABC-F proteins is highly conserved [23, 24]. Valnemulin-ribosome docked compounds revealed that the tricyclic mutilin core on the base of C2452, U2505, U2506, A2062, G2061, and its C14 extension was stabilized by U2585 [2, 13, 14]. As the SrpA loop residues Met218, Met219, and Trp213 were in direct contact with the crucial nucleotides A2062, G2061, U2585, and C2452 surrounding the valnemulin molecule, the insertion of the SrpA loop region might cause conformational changes of the valnemulin-binding pocket. Furthermore, Met219 of SrpA directly competed with valnemulin, similar to Leu242 in MsrE.

The linker connecting the two tandem ABC domains is a defining feature of the ABC-F family, and the loop region of the linker forms stereospecific contact near the PTC in the ribosome [20]. Unlike MsrE and VgaA_LC_ variants, truncation of the loop region completely abolished resistance to pleuromutilins, lincosamides, and streptogramin A [22, 23]. Furthermore, Lys206Ala, Ser211Ala, Trp213Ala, Met218Ala, and Met219Ala substitutions reduced PLS_A_ resistance to different degrees. This variation is presumably because mutations reduce the size of the sidechain that penetrates most deeply into the PTC in SrpA, thereby eliminating steric clashes with valnemulin and reducing the magnitude of the relatively small allosteric conformational changes in the PTC upon SrpA binding [20].

The mechanism by which ABC-F proteins mediate antibiotic resistance has been a subject of long-standing controversy. The efflux hypothesis posits that ABC-F proteins export antibiotics out of the cell, similar to other ABC superfamily members, while a recent study has shown that ABC-F proteins mediate antibiotic resistance by interacting with the ribosome and displacing the bound drug [20, 25]. In this study, we observed that the addition of the putative efflux pump inhibitor carbonyl cyanide m-chlorophenylhydrazone (CCCP) (10 μg/mL) or reserpine (20 μg/mL) had no effect on PLS_A_ resistance in isolates harboring *srpA* (data not shown). A ribosome binding assay unambiguously demonstrated that SrpA can prevent that binding of antibiotics to ribosomes and displaces ribosome-bound antibiotics.

A BLASTp search against the NCBI database revealed that SrpA is exclusively found in *S. suis*, with a number of SrpA-like proteins (82–99% amino-acid identity) distributed across isolates from China, USA, UK, Vietnam, and the Netherlands, indicating the global dissemination of SrpA. Interestingly, in most isolates, such as 05ZYH33 (CP000407.1), 98HAH33 (CP000408.1), and P1/7 (AM946016.1), *srpA* could be divided into three adjacent ORFs separated by 100–200 bp, respectively. The first ORF (51 amino acid) showed 88% amino acid identity to SrpA (residues 1– 16), the second ORF (180 amino acid) showed 89% identity (residues 34–212), while the last ORF (249 amino acid) showed 81% identity (residues 218–455). We speculate that a loop truncation explains the pleuromutilin susceptibility in these strains.

In the present study, all *srpA* genes were located on chromosomes and were intact. With respect to genetic context, most of the isolates belonged to type I (n = 40), in which *srpA* was invariably located between *pbp2b* (WP_105203009.1) and *yycJ* (WP_024376691). However, isolates without *srpA* harbored a similar arrangement, separated by 400–500 bp. Only some antibiotic resistance ABC-F proteins mediating the PLS_A_ resistance phenotype have been found to be chromosomally encoded and embedded in genes, including *vmlR* in *B. subtilis, lsa*(A) in *E. faecalis, tva*(A) in *Brachyspira hyodysenteriae, lsa*(A) in and *salA* in *S. sciuri* [38, 47-49]. The genetic context of type II isolates was highly divergent from that of type I isolates, with two directly repeated IS481 family IS*Ssu8-*like elements in the flanking region of *srpA*; this segment could form a circular intermediate composed of *srpA* and an IS*Ssu8-*like element. Since *S. suis* has been recognized as a reservoir for the spread of resistance genes to major streptococcal pathogens, the potential risk of disseminating of *SrpA* from *S. suis* to other *Streptococcus* spp. is worrisome [27, 50]. The closest NCBI matches to the type II segment were regions in *S. suis* 1081 and *S. suis* 0061 from China (CP017667.1 and CP017666.1) (71% amino acid identity), but the IS*Ssu8-*like elements were absent from both. In light of these findings, we speculated that *srpA* might stem from *S. suis* and confers innate resistance to PLS_A_. It is conceivable that over evolutionary timescales, some *S. suis* isolates stably inherited the resistance gene from a common ancestor, while other isolates lost the ancestral *srpA* gene. Given that is *S. suis* currently exposed to numerous PLS_A_ antibiotics in veterinary clinics, the appearance of *srpA* on mobile genetic elements is a feasible survival strategy in PLS_A_-susceptible *S. suis*.

In summary, in this study, a new antibiotic resistance ABC-F family determinant, *srpA*, was detected; the gene confers cross-resistance to pleuromutilins, lincosamides, and streptogramin A in *S. suis* and *S. aureus*. SrpA possessed the characteristic ABC-F family protein conformation and showed the highest similarity with Vga(E). Molecular docking suggested that the antibiotic resistance domain loop region of SrpA penetrated deep into the PTC cavity and occupied the valnemulin-binding pocket. Mutagenic studies implicated the ARD loop region in mediating the specificity and efficiency of antibiotic resistance. Furthermore, ATP hydrolysis residues (103E/408E) played a paramount role in SrpA activity. Importantly, we found that SrpA can protect ribosomes and promote the dissociation of the drug from its binding sites, suggesting that a similar ribosomal protection mechanism is shared by antibiotic resistance ABC-F family proteins.

## Abbreviations

MDR: Multidrug-resistant
MRSA: methicillin-resistant Staphylococcus aureus
VRE: vancomycin-resistant Enterococcus faecium
PTC: peptidyl-transferase center
TMDs: transmembrane domains
VAL: valnemulin
ENR: enrofloxacin
FPIA: fluorescence polarization immunoassay
NBSs: nucleotide-binding sites
FP: fluorescence polarization
CCCP: carbonyl cyanide m-chlorophenylhydrazone

## Acknowledgements

We are grateful for the sampling support from microbiologists in these agencies: Chongqing Academy of Animal Sciences, Sichuan Provincial Agricultural Department, Sichuan Agricultural University, Qingdao Agricultural University and Henan Agricultural University.

## Conflicts of interest

No potential conflict of interest was reported by the authors.

**Fig. S1.**
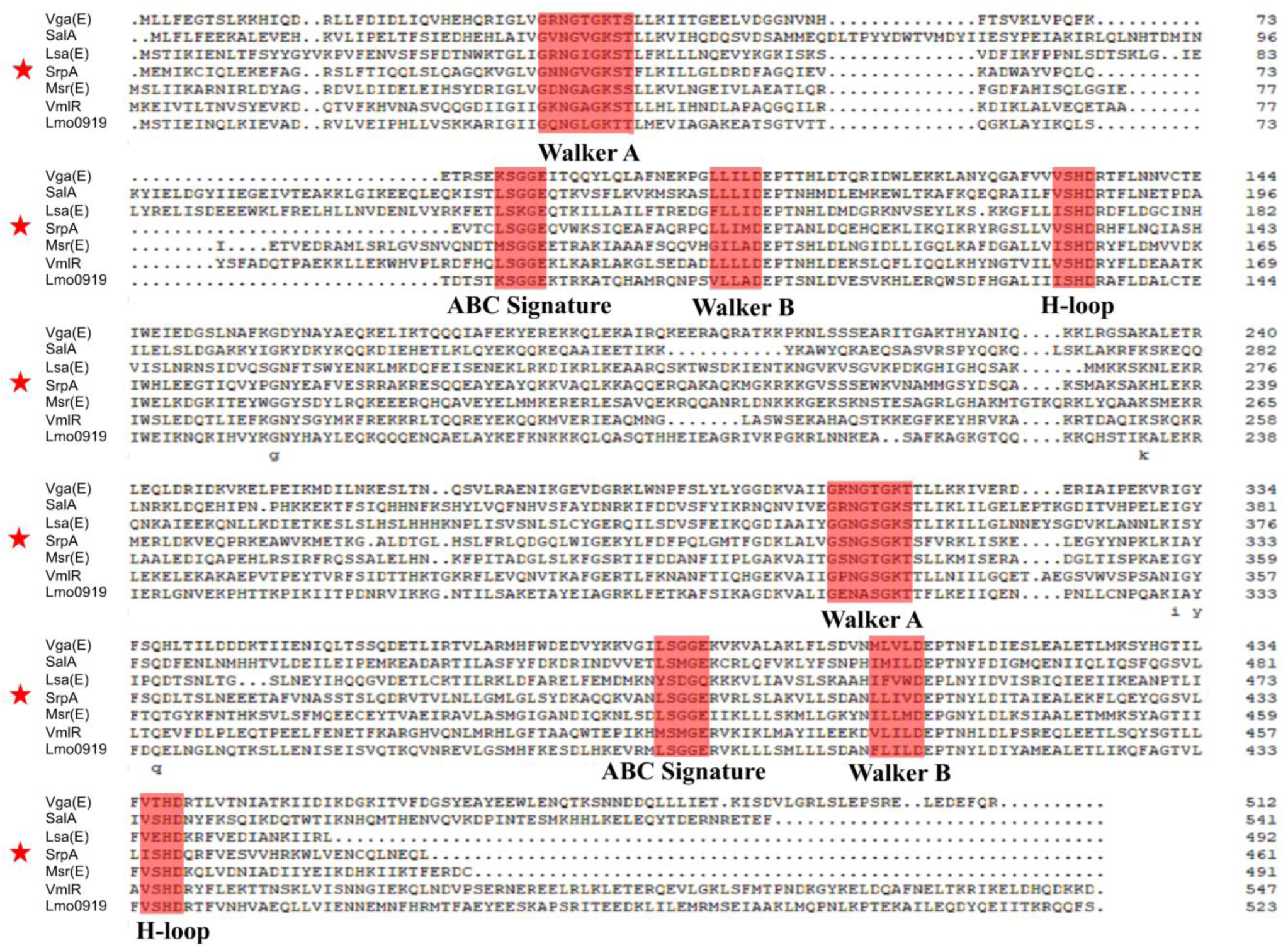
Multiple alignment of SrpA-related proteins. Analysis was carried out with Sal(A), Lsa(E), Vga(E), Msr(E), VmlR, and Lmo0919. Two copies of the Walker A and Walker B motifs as well as ABC signatures and H-loops are indicated in red, and SrpA is indicated by an asterisk.

**Figure. S2.**
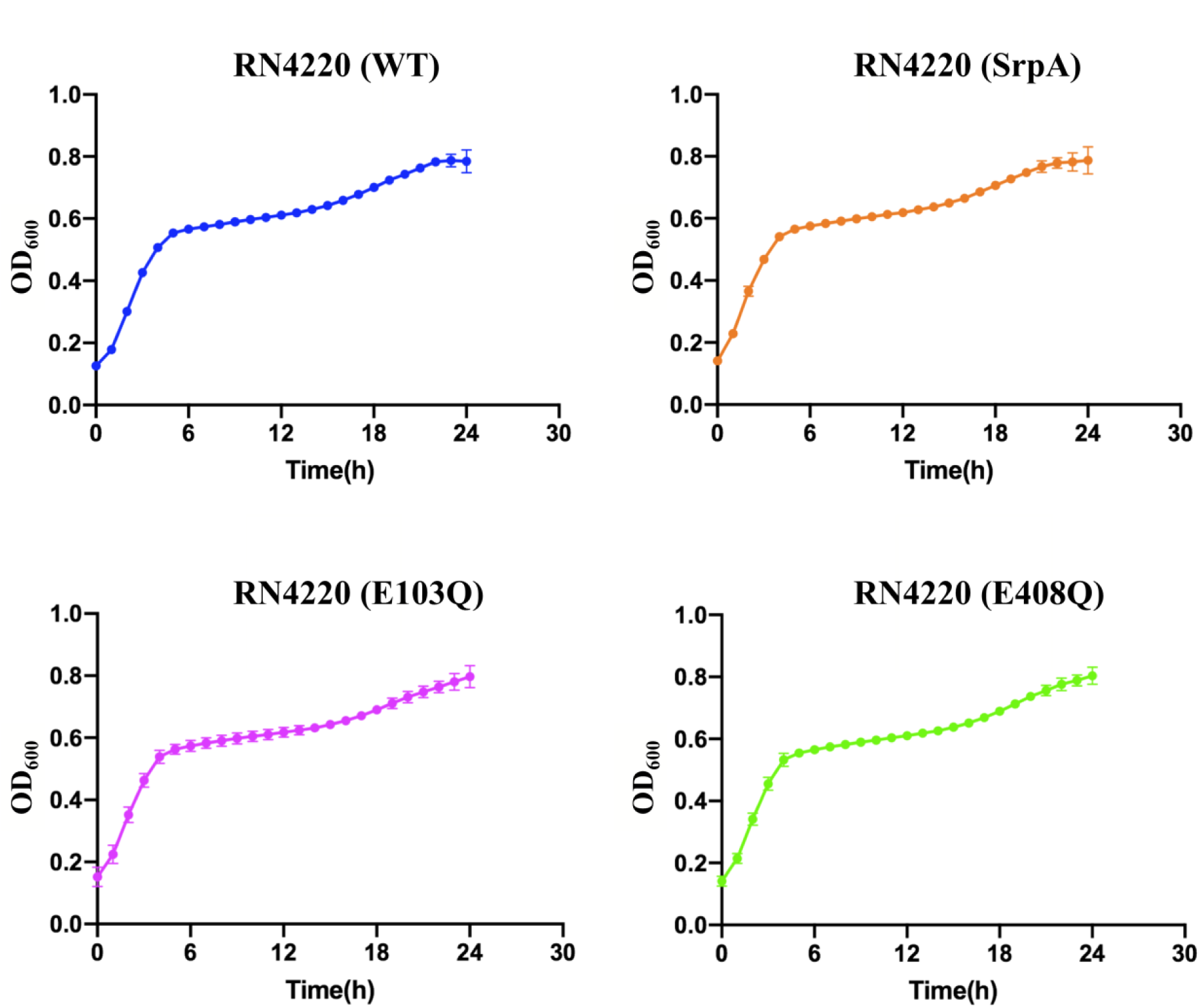
Effect of SrpA expression on *S. aureus* growth. Wild-type *S. aureus* RN4220 was used as a control (blue), and the RN4220/pAM401-SrpA, RN4220/pAM401-SrpA(E103Q), and RN4220/pAM401-SrpA(E408Q) are indicated in orange, magenta, and green, respectively.

**Fig. S3.**
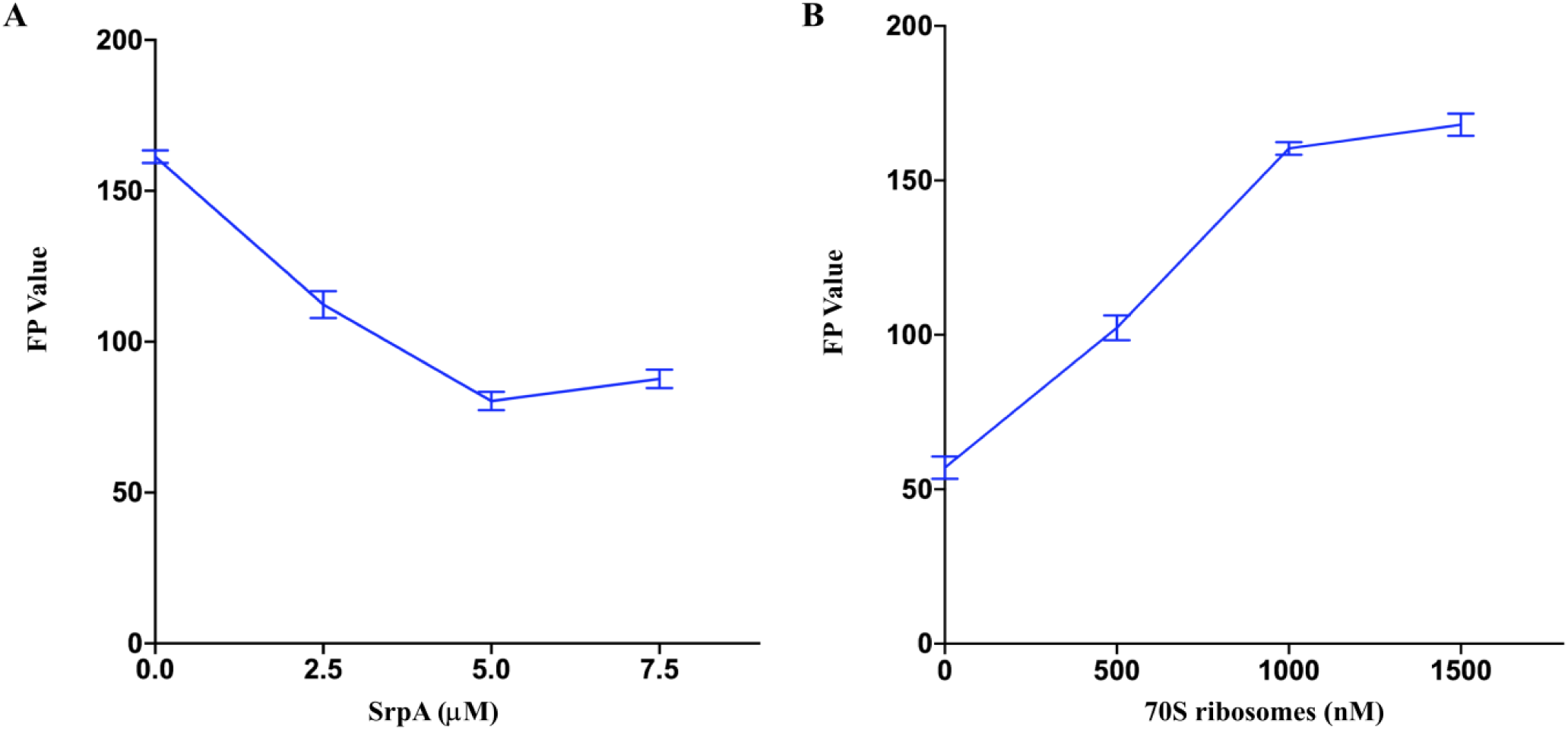
Optimization of substrate concentrations to assess the stoichiometry of the SrpA binding assay. (A) Concentration of 70S ribosomes. (B) Concentration of SrpA. Results are means from at least three independent determinations, and error bars show standard deviations.

**Figure. S4.**
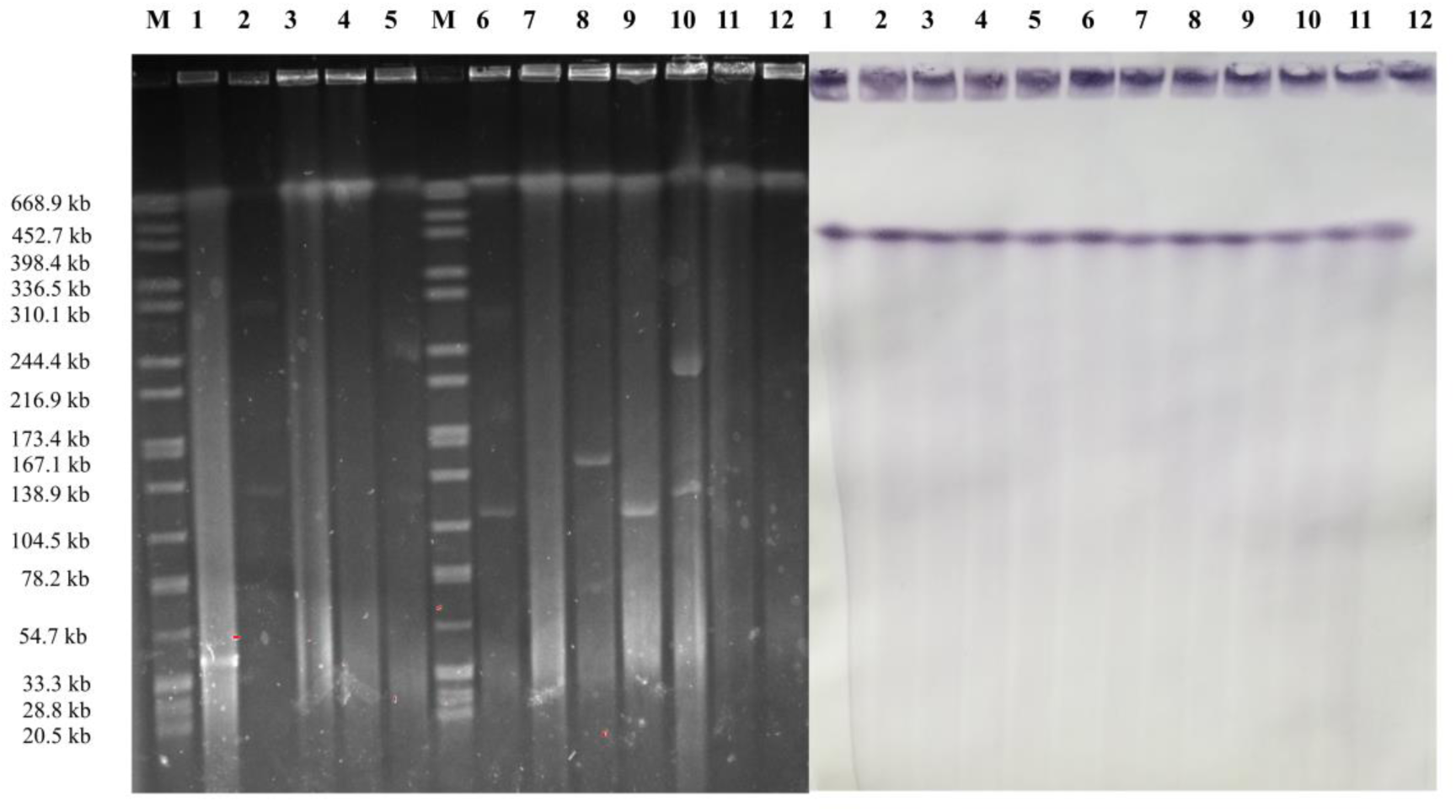
S1-PFGE analysis of *srpA-*positive *S. suis*. Bands M: H9812, 1: HNBY78, 2: HNBY23, 3: SD2B12, 4: SD2B15, 5: CQ2B66, 6: F5-1HN, 7: BJAY75, 8: BJCY31, 9: SC5B43, 10: CQ1A4, 11: SC5B66, 12: LZ11G.

**Table. S1.**
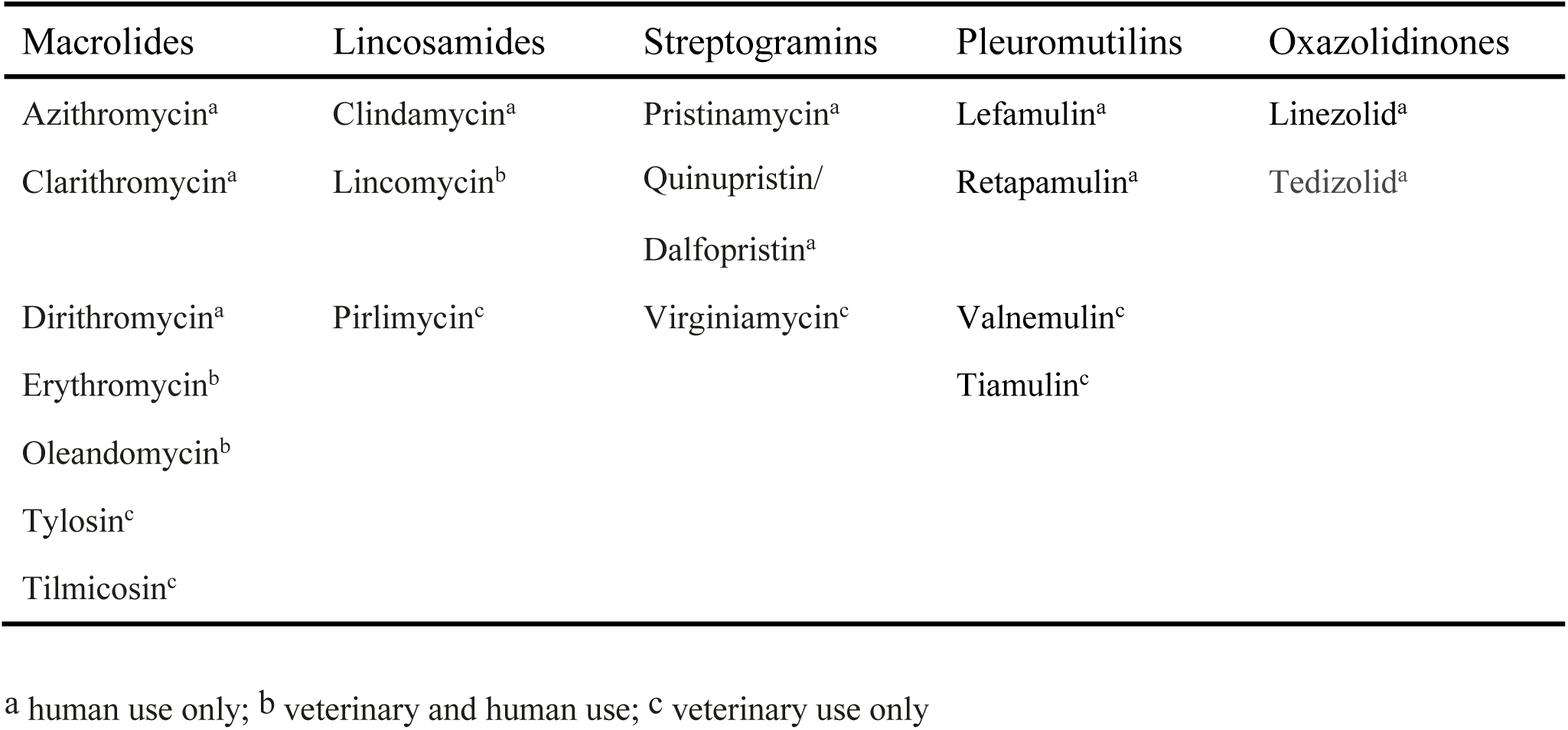
List of commonly used macrolides, lincosamides, pleuromutilins, oxazolidinones and streptogramins for human or veterinary.

| Macrolides | Lincosamides | Streptogramins | Pleuromutilins | Oxazolidinones |
| --- | --- | --- | --- | --- |
| Azithromycin<sup>a</sup> | Clindamycin<sup>a</sup> | Pristinamycin<sup>a</sup> | Lefamulin<sup>a</sup> | Linezolid<sup>a</sup> |
| Clarithromycin<sup>a</sup> | Lincomycin<sup>b</sup> | Quinupristin/<br>Dalfopristin<sup>a</sup> | Retapamulin<sup>a</sup> | Tedizolid<sup>a</sup> |
| Dirithromycin<sup>a</sup> | Pirlimycin<sup>c</sup> | Virginiamycin<sup>c</sup> | Valnemulin<sup>c</sup> |  |
| Erythromycin<sup>b</sup> |  |  | Tiamulin<sup>c</sup> |  |
| Oleandomycin<sup>b</sup> |  |  |  |  |
| Tylosin<sup>c</sup> |  |  |  |  |
| Tilmicosin<sup>c</sup> |  |  |  |  |

<sup>a</sup> human use only; <sup>b</sup> veterinary and human use; <sup>c</sup> veterinary use only

**Table. S2.**
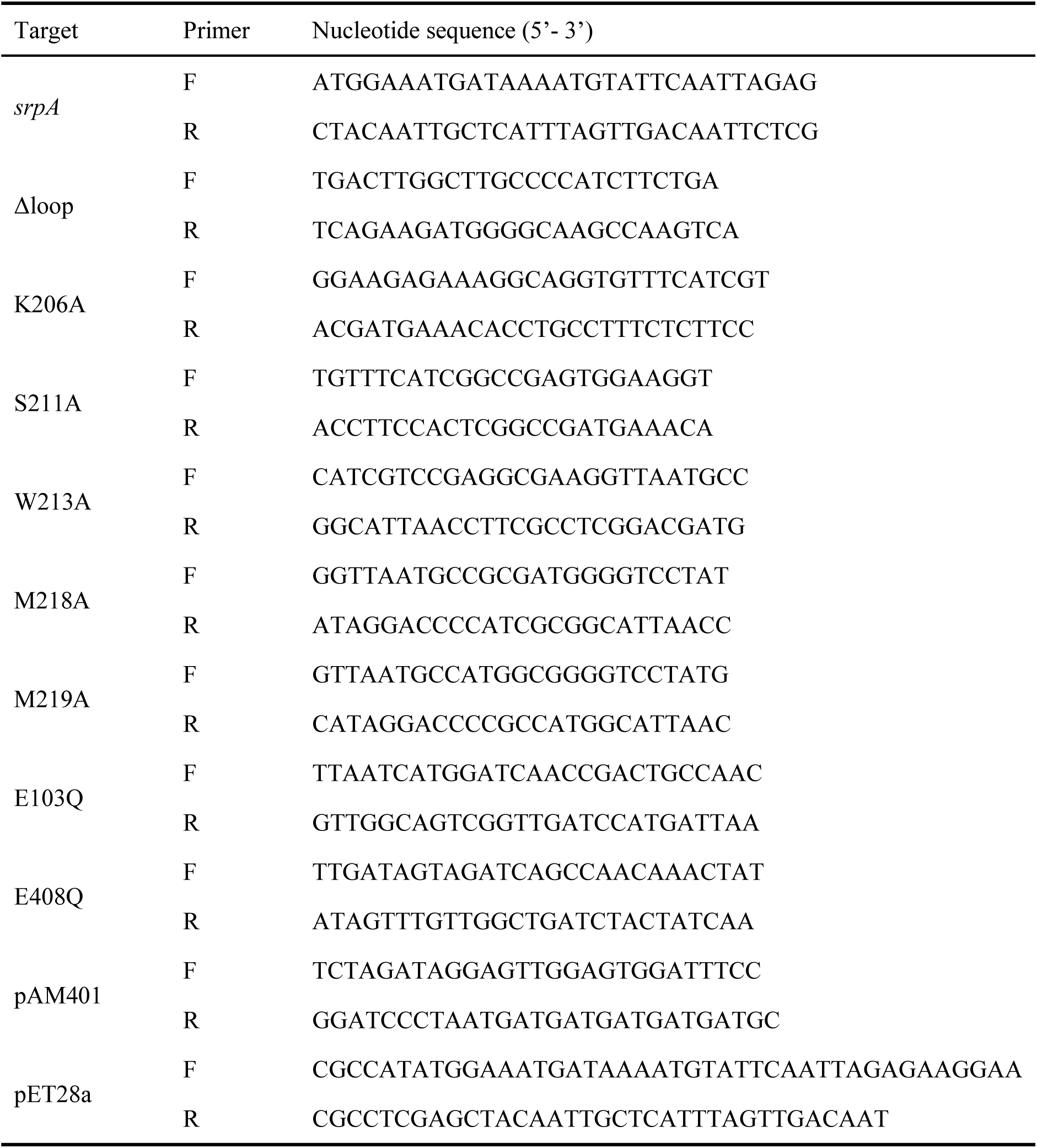
Primers used in this work

